# coda4microbiome: compositional data analysis for microbiome studies

**DOI:** 10.1101/2022.06.09.495511

**Authors:** M.Luz Calle, Antoni Susin

## Abstract

**Motivation:** One of the main challenges of microbiome analysis is its compositional nature that if ignored can lead to spurious results. This is especially critical when dealing with microbiome variable selection since classical differential abundance tests are known to provide large false positive rates.

**Results:** We developed coda4microbiome, a new R package for analyzing microbiome data within the Compositional Data Analysis (CoDA) framework in both, cross-sectional and longitudinal studies. The core functions of the library are aimed at the identification of microbial signatures and involve variable selection in generalized linear models with compositional covariates. All algorithms are accompanied by meaningful graphical representations that allow a better interpretation of the results.

**Availability:** coda4microbiome is implemented as an R package and is available at CRAN https://cran.r-project.org/web/packages/coda4microbiome/index.html.

**Supplementary information:** coda4microbiome project website: https://malucalle.github.io/coda4mi-crobiome/.

## 1 Introduction

Although there are still many unknowns about the specific mechanisms of action of the human microbiome, there is growing evidence of its relevance in human health (Lo et al. 2021, Zheng et al. 2020). In recent years, much progress has been made in microbiome research thanks to high-throughput DNA sequencing technologies that allow precise quantification of the composition of the microbiome. The study of the microbiome is considered a great opportunity for improving the current treatment of some diseases and for deriving microbial biomarkers that could be used as diagnostic or prognostic tools.

The analysis of microbiome data involves significant experimental and computational challenges (Bharti and Grimm, 2021). One of them is the compositional nature of the data, which requires the use of specific methods of analysis (Calle 2019, Gloor et al. 2017). Compositional data refers to constraint multivariate non-negative data that carry relative information. Microbiome relative abundances (proportions) are constrained by a total sum equal to one. This total constraint induces strong dependencies among the observed abundances of the different taxa. In fact, the observed abundance of each taxon is not informative and only provide a relative measure of abundance when compared to the abundances of other taxa (Susin et al. 2020). Ignoring the compositional nature of microbiome data can lead to spurious results. This is especially critical when dealing with microbiome variable selection. In this context, several simulation studies have shown that commonly used differential abundance tests provide large false positive rates (Weiss et al. 2017, Nearing et al. 2022).

Aitchison (1982) laid the foundations of Compositional Data Analysis (CoDA), which relies on extracting the relative information of compositional data by comparing the parts of the composition. Logarithms of ratios between components (log-ratios) are the fundamental transformation in this framework (Pawlowsky-Glahn et al. 2015, Greenacre 2021). The applicability of CoDA methods in microbiome studies has been hindered by the availability of software and the difficulty of interpreting their results.

Here we introduce coda4microbiome, a new R package for analyzing microbiome data within the CoDA framework in both, cross-sectional and longitudinal studies. The core functions of the library are aimed at the identification of microbial signatures and involve variable selection in generalized linear models with compositional covariates. All algorithms are accompanied by meaningful graphical representations that allow a better interpretation of the results.

In fact, coda4microbiome is not just an R package but a broader initiative that aims to bridge the gap between compositional data analysis and microbiome research. To this end, we are conducting training activities and developing materials that are available at the website of the project: https://malucalle.github.io/coda4microbiome/.

## 2 Materials and Methods

The algorithms implemented in coda4microbiome package rely on the analysis of log-ratios between components. Here we describe the analysis for cross-sectional microbiome data. The algorithms for longitudinal microbiome data are described in Calle and Susin (2022).

### 2.1 Microbiome signature based on log-ratio analysis

Assume we have n subjects with phenotype *Y* = (*Y*_1_, …, *Y*_*n*_) and denote by *X*_*i*_ = (*X*_*i*1_, *X*_*i*2_, *…, X*_*iK*_) the microbiome composition of subject i for K taxa. *X* can represent either relative abundances (proportions) or raw read counts. The identification of those taxa that are associated to the outcome can be approached through penalized regression on a generalized linear model with a zero-sum constraint on the regression coefficients (Lu et al. 2018):

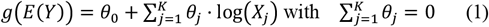

This is known as a log-contrast model (Aitchison and Bacon-Shone, 1984) and the zero-sum constraint ensures the scale invariance principle required in CoDA.

Model (1) can be reparametrized and expressed as the “all-pairs log-ratio model”, a generalized linear model containing all possible pairwise logratios (Bates and Tibshirani, 2018):

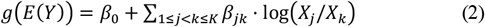

Our algorithm performs variable selection through penalized regression on Model (2). The regression coefficients in equation (2) are estimated to minimize a loss function *L*(*β*) subject to an elastic-net penalization term on the regression coefficients:

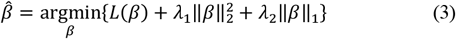

A common reparameterization of the penalization parameters is *λ*_1_ = *λ*(1 − α)/2 and *λ*_2_ = *λ*α where *λ* controls the amount of penalization and α the mixing between the two norms (Friedman et al. 2010).

For the linear regression model the loss function is given by the residual sum of squares 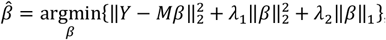, where *M* is the matrix of all pairwise log-ratios and has dimension n by *K*(*K* − 1)/2. The expression of the optimization problem (3) for other models, like the logistic regression, can be found in Friedman et al. (2010). We use the function cv.glmnet() from the R package glmnet (Friedman et al. 2010) to solve (3) within a cross-validation process that provides the optimal value of *λ* with a default value for α equal to 0.9.

The result of the penalized optimization provides a set of selected pairs of taxa, those with a non-null estimated coefficient. For each individual *i*, the linear predictor of model (2) provides its microbiome signature score, given by 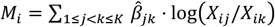. Because of the linearity of the logarithm, this microbiome signature *M* can be rewritten in terms of the selected single taxa which is more interpretable than the selected pairs of components:

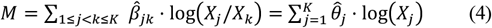

where 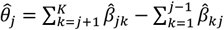 (Bates and Tibshirani, 2018).

It can be proved that 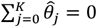 and thus, the microbiome signature *M* is a log-contrast function involving the selected taxa (those with 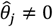). This ensures the invariance principle required for proper compositional data analysis and it facilitates the interpretation of the microbiome signature: Expression 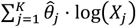 in (4) can be interpreted as a weighted balance between two groups of taxa, *G*_1_ and *G*_2_, the taxa with a positive coefficient vs those with a negative coefficient (Susin et al. 2020).

The results of the analysis are visualized with a plot of the microbial signature where the selected taxa and the corresponding coefficients are represented in a bar plot (Fig. 1). A second plot describes the discrimination capacity of the selected microbial signature.

**Fig. 1.**
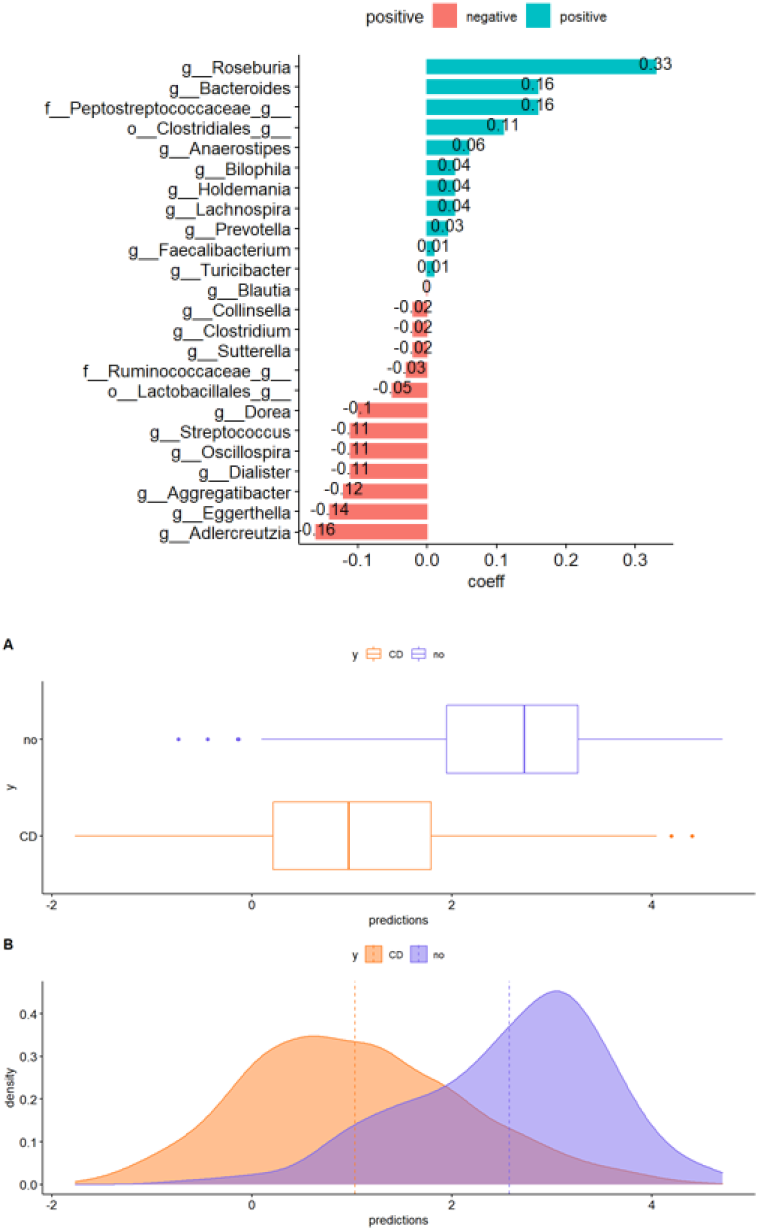
Example of coda4microbiome plots for Crohn’s disease (Rivera-Pinto et al. 2018). Top: microbial signature bar plot (green: positive coefficients and red: negative coefficients). Below: distribution of the microbial signature scores, boxplot (A) and density plot (B) (orange: Crohn’s disease, blue: controls)

### 2.2 coda4microbiome main functions

#### Functions for microbial signature identification

- **coda_glmnet**: Identification of microbial signatures in cross-sectional studies. This function performs variable selection through penalized regression on the set of all pairwise log-ratios for both, binary outcome (logistic regression) and continuous outcome (linear regression). The result is expressed as the (weighted) balance between two groups of taxa. It allows the use of non-compositional covariates. A graphical representation of the taxa that constitutes the signature and their coefficients is provided. The output also includes a plot of the prediction or classification accuracy.
- **coda_glmnet_longitudinal**: Identification of microbial signatures in longitudinal studies. This function identifies a set of microbial taxa whose joint dynamics is associated with the phenotype of interest (binary). The algorithm performs variable selection through penalized regression over the summary of the log-ratio trajectories (AUC). The result is expressed as the (weighted) balance between two groups of taxa. The output provides three plots: the taxa that constitutes the signature and their coefficients, the classification accuracy of the signature and the plot of the signature trajectories of the individuals.

#### Functions for log-ratio exploratory analysis

Previously or independently of variable selection for microbial signature identification, one may be interested in the exploratory analysis of pairwise log-ratios. However, the interpretation of results of pairwise log-ratio analysis is challenging because when one taxon A is highly associated with the outcome, any log-ratio involving taxon A is likely to be associated with Y, no matter which is the second taxon involved in the logratio. The following two functions summarize the importance of each taxon A by aggregating the prediction accuracy of all pairwise log-ratios that involve taxon A. The output includes a correlation-like plot of the association of each pairwise log-ratio with the outcome where the taxa are ranked by importance.

- **explore_logratios**: Explores the association of each log-ratio with the outcome.
- **explore_lr_longitudinal**: Explores the association of a summary (integral) of each log-ratio trajectory with the outcome.

#### Suplementary functions

- **explore_zeros**: Provides the proportion of zeros for a pair of variables (taxa) and the proportion of samples with zero in both variables. A bar plot with this information is also provided. Results can be stratified by a categorical variable.
- **impute_zeros**: Simple imputation: When the abundance table contains zeros, a positive value is added to all the values in the table. It adds 1 when the minimum of table is larger or equal to 1 (i.e. tables of counts) or it adds half of the minimum value of the table, otherwise.
- **logratios_matrix**: Computes the matrix with of all pairwise log-ratios between taxa.
- **plot_prediction**: Plot of the predictions of a fitted model (microbial signature): Multiple box-plot and density plots for binary outcomes and regression plot for continuous outcome.
- **plot_signature**: Graphical representation of the variables selected and their coefficients.
- **coda_glmnet_null**: Performs a permutational test for the coda_glm_net() algorithm. It provides the distribution of results under the null hypothesis by implementing the coda_glmnet() on different random rearrangements of the response variable.
- **filter_longitudinal**: Filters those individuals and taxa with enough longitudinal information.
- **coda_glmnet_longitudinal_null**: Performs a permutational test for the coda_glmnet_longitudinal() algorithm. It provides the distribution of results under the null hypothesis by implementing the coda_glmnet_longitudinal() on different random rearrangements of the response variable.
- **shannon, shannon_effnum and shannon_sim**: Shannon information, Shannon effective number of variables in a composition and Shannon similarity between two compositions, respectively.

## 3 Conclusions

We developed an R package for microbiome analysis that deals with the compositional nature of microbiome data. coda4microbiome provides a set of functions to explore and study microbiome data within the CoDA framework, with a special focus on identification of microbial signatures that can serve as biomarkers of disease risk and prognostic. The results are expressed as the (weighted) balance between two groups of taxa, those that contribute positively to the microbial signature and those that contribute negatively.

With this new R package we aim to enhance microbiome analysis by taking into consideration the compositional nature of microbiome data through the use of compositional data analysis methods. The algorithm is implemented in the R package “code4microbiome” (https://cran.r-project.org/web/packages/coda4microbiome/) that is accompanied with a vignette with a detailed description of the functions. The website of the project contains several tutorials: https://malucalle.github.io/coda4micro-biome/.

